# Examining Anxiety and Risk-taking in Healthy Male and Female Wistar Rats using Spatial and Temporal Analysis of Elevated Plus Maze

**DOI:** 10.1101/2022.11.16.516842

**Authors:** Sakshi Sharma, Jyotsna Pandey, Suman Jain, Varsha Singh

## Abstract

The Elevated Plus Maze (EPM) offers a standard set up for understanding anxiety, unconditioned risk-avoidance in rodents. The animal shows a preference for enclosed safe arms and avoids risky, open arms that evoke anxiety due to elevated platform (unconditioned response to elevation). A few rodent studies aiming to understand sex-skewed representation in anxiety disorders use the maze and report that more females compared to males show high levels of anxiety and risk-avoidance on an elevated platform. Ethograms derived from examining animal behaviour in the EPM provide precise measures of behaviour of interest, enabling objective assessment of anxiety and risk behaviour. We report two new parameters that might be critical for quantitative analysis of EPM task as a measure of anxiety with reference to sex-differences in risk-taking: (a) spatial preference for arms (open & closed) (b) temporal shift in arm preference in task trials. We first report results from the conventionally derived measures confirming that males spent more time in the open arms indicating low anxiety and higher risk-taking. Preferences for the two open and two closed arms were non-uniform for males and females; male showed stronger preference for one of the two risky open arms, and females showed a stronger preference for one of the two safe closed arms. Temporal analyses indicated that males spent more time in open arms in 3 out of 6 time bins (time blocks), and females spent more time in the closed arms in 5 out of 6 time bins (time blocks), however, counter-intuitively, females showed larger increase in time spent in open arms in the last phase potentially indicating greater regulation of anxiety and increased risk-taking. Inclusion of spatial and temporal parameters in EPM studies might improve our understanding of cognitive and biological sex-differences pertaining to anxiety, and risky behaviour.

## Introduction

The elevated plus maze (EPM) is a measure of risk and exploration, used for understanding effects of interventions that help overcome anxiety in rodents. The maze consists of the four arms perpendicular to each other, two are closed and the other two are open and because the maze is kept at a height, the elevation of open arms with no protective walls serves as a natural stressor and is avoided whereas the closed arms provide safer environment to the animal. The maze makes use of rodents’ unconditioned preference for safe, dark, closed spaces and an inherent fear of risky, open spaces and heights to provide an effective laboratory-based measure of anxiety (Barnett, 1975). The animal tends to spend more in the closed arms reflecting avoidance of risk, seeking safety/protection due to high levels of anxiety, anxiety-reducing intervention increases the time spent in the open arms indicative of low levels of anxiety, higher exploration and risk taking (Korte, 2003; Yamaguchi, 2005; Carobrez 2005; Walf, 2007; Pinheiro, 2007; Campos, 2013).

A standard experiment using EPM consists of a single trial of 5 minutes duration where the animal is kept in the center of the maze and is allowed to explore the four arms for 5 minutes. Although a widely used measure, EPM has been analysed in different ways, for instance, apart from time spent in the open or closed arms, behaviour that disrupts animal’s physical/postural stability such as incomplete entries to the four arms, head dips and rearing (Wall, 2001), indicates high levels of anxiety and risk-avoidance (Filgueiras et al, 2014). Numerous modifications of the standard trial on EPM have been used over the years to investigate the changes in the animal’s behavior on EPM if exposed to different experimental designs. Handley and Mithani (1984) examined anxiety levels by assessing the ratio of open arm entries to total arm entries. Pellow (1984) examined anxiety by using the ratio of time spent in each arm. Time spent in the open arms is a commonly used measure of anxious phenotype in the EPM (Silva et al, 2013; Weinstock, 2016; Coimbra et al, 2017; Himanshu; 2020).

One problem in the EPM studies that rely on time spent in open versus closed arm and arm-preference is that it discounts non-uniformity of two open arms (open sub-arm 1 and 2) and two closed arms (closed sub-arm 1 and 2), preference for a one arm/side has been observed (Schwarting, 2005). Another problem in the EPM studies is that the avoidance of open arm as a measure of unconditioned anxiety and risk avoidance due to a single task exposure causes an emotion-shift; “one-trial tolerance” that is an earlier exposure to EPM (trial 1) diminishes anxiety-reducing effect of pharmacological/anxiolytic effect on trial 2 (File, Mabbutt, and Hitchcott, 1990). This non-pharmacological exposure of trial 1 reducing the effects of anti-anxiety intervention on trial 2 is attributed to cognitive-affective learning that the open arms will not lead to escape, thereby reducing exploration of open arm and reducing the approach-avoidance conflict related to open arm (Rodgers & Sheperd, 1993). File et.al (1995) used two trials of different durations, to study the effect of anxiolytic drug, diazepam. They reported that diazepam had significant anxiolytic effect for both the 5min and 10 min trial duration, however, second trial for 5 min duration did not show effects of diazepam, which were sustained in the second trial for 10 min duration. The timepoint in the EPM trial (5 mins) where the shift in anxiety and open arm preference occurs remains unclear, however, temporal analysis suggests that anxiolytic effects might not sustain beyond 3 mins (Rosa et al., 2000).

Although EPM is used for assessing anxiety and risk-taking in healthy male and female Wistar rats (Walf and Frye, 2007), inconsistencies in sex-specific anxiety and risk taking on the EPM (Elliott et al., 2004; Genn et al., 2003; Scholl et al., 2019; Winther et al., 2018; Mansouri et al., 2019; Yang et al., 2019; Mansouri et al., 2019; Sakhaie et al., 2020) might be due to discounting of sex-specific impact of spatial and temporal features in EPM analysis. Studies have also pointed out that temporal and spatial features tend to impact anxiety levels on EPM (Schneider, P., Ho, Y. J., Spanagel, R., & Pawlak, C. R., 2011). Therefore, we expect that detailed analysis of spatial and temporal aspects of EPM might improve our understanding of sex-differences in anxiety, for instance, modified EPM (elevated gradient of aversion, EGA) using shortened trial duration to 3 min indicated female rats might be less anxious compared to male counterparts (Bonuti, 2022). Donner and Lowry, 2013; Pavlova et. al, 2020; Knight et.al, 2021 have also reported similar differences in the two sexes with females showing less anxious behavior on the maze. We expected that examining sex-specific spatial variation in arm preference, and variability due to temporal feature of EPM might potentially improve our understanding of sex-differences in anxiety and risk taking.

## Methods

### Animal

Adult male (n=10) and female (n=10) albino Wistar rats weighing 240-280g were used for this study. Animals were housed in separate polypropylene cages and provided with food pellets and water ad libitum. A room temperature of 25°C and light: dark cycle of 10:14h were maintained. All experiments were conducted under the norms set up by Institutional Animal Ethics Committee.

### Sample size and selection of subject

The sample size estimation was done by ‘Resource Equation Method’. According to this method, degree of freedom of ANOVA is calculated, which is termed as “E” value. E value is measured by the mathematical formula, E = Total number of animals – Total number of groups. The value between 10 to 20 is considered as an adequate value and is accepted.

### Elevated Plus Maze (EPM)

We made use of EPM setup composed of plus maze built from opaque plastic board attached to iron and wooden frame. The dimensions for the open arms were 76cm in length and that of the closed arms were 134cm. The center area was 6×6cm in its dimensions. Video-tracking of behavior of the animal was done with CCD camera (Panasonic, Japan).

Single day protocol was performed with prior habituation of the animal to the testing room where the experiment was performed for 30minutes. This was followed by placing of the animals in the central area of the maze facing any of the two arms of the open area. This marked the start of a trial which lasted for 5minutes where animals were allowed to explore in the maze.

### Spatial and Temporal Features of EPM

We examined spatial and temporal features in EPM to understand male-female differences in anxiety and risk-taking. Behavioral tags were obtained from a widely used software, Kinoscope that produces output in the form of a visual map (Kokras et al., 2017), it served as inputs for manual scoring of male-female rodent behaviour in the two open versus closed arms of EPM. Simultaneous scoring of the behavioural parameters was done by the software and manual scoring was done for the head dips and rearing. The software tags behaviour (like protected or unprotected head dips) which is followed by the manual scoring for the duration of 300s (trial duration for EPM). The visual maps represented events/ behaviour of interest from the start to the end of a trial, i.e., from 0^th^ to 300^th^ second (colour coded, pl. see Fig1.2). Time spent in the two open and closed arms and other regions of the maze was examined. Events obtained were a) incomplete entries to each of the specific areas of the area i.e., closed arms, open arms and the centre area and b) number of protected and unprotected head dips in both the groups (c) time spent in each of the areas within the bin of 50s to study the difference in behaviour with course of time.

### Statistical Analysis

Shapiro-Wilk normality test was performed for the distribution of data. Sample size was estimated by Resource Equation Method (Festing, 2002) used as per previous reports. 1) Time spent in the zones of arena, namely Open arms and Closed arms along with the ratio of time spent in the Open arms to that of Closed arms and 2) Chronology of the time spent in these areas within a bin of 50s were tested between the two groups, i.e., males and females. T-test was performed for the comparison within the two groups (non-parametric, two-tailed and unpaired). Repeated measured ANOVA was performed to check the differences between the means of the time spent in the 6 bins examined. Visual maps or ethograms were obtained by the algorithm used by Kinoscope software. Data was represented as individual values of mean and standard error of mean and the p-value <0.05 was accepted as significant.

## Results

### 1. Spatial Feature: Arm Preference

#### 1.1 Ratio of time spent in Open to Closed arms

When the ratio of the time spent in the open arms to that of closed arms was measured (Fig 1.1A), it was found to be greater in case of the males (t=3.3; p=0.002; Cohen’s d=1.475; df=9). In case of the time spent on the maze (Fig1.1B), males (mean =62.38s) were observed to spend more time in the open arms than the females (mean=28.48s) with the significant difference (t=2.87; p=0.005; Cohen’s d= 1.29; df=9) and consequently females (mean=199.8s) spent more time in the closed arms than the males (mean=137.68s) with the significant difference (t=−3.87; p=0.001; Cohen’s d=1.73; df=9).

**Figure 1.1.**
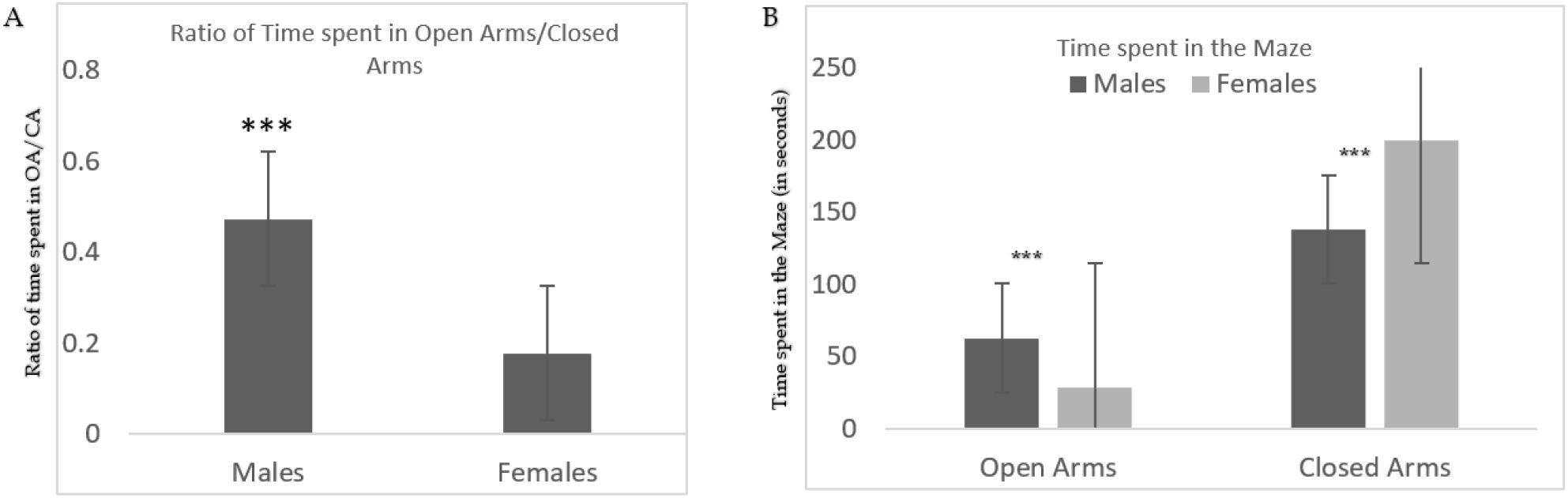
Increased anxiety-like behaviour exhibited by the males compared to the female rats on the elevated plus maze test. Figure shows the ratio of time spent in the open arms to that of the closed arms (1.1A) and the mean time spent in the open and closed arms of the elevated plus maze by the males and the females (1.1B). Males spent more time in the open arms and displayed less anxiety-like behaviour. N=10/group. Data expressed as means ± SEM. (***: p-value<0.05).

#### 1.2. Non-uniform Arm Preference

The colour coded visual maps by Kinoscope show the increased number of behavioural events by the males on the EPM when compared to the females (Fig1.2A and 1.2B). We observe that the time spent in the open arms (marked with red and black) was more by the males than females. 90% of the males show increased behavioural events with more time spent in the open arms whereas, 70% of the females show decreased events with more time spent in the closed arms (marked with blue and yellow). Behavioural events like incomplete entries to the open (red bars) and closed arms (marked with black bars) and head dips {both protected (yellow bars) and unprotected (green bars)} were also exhibited more by males than the females.

**Figure 1.2.**
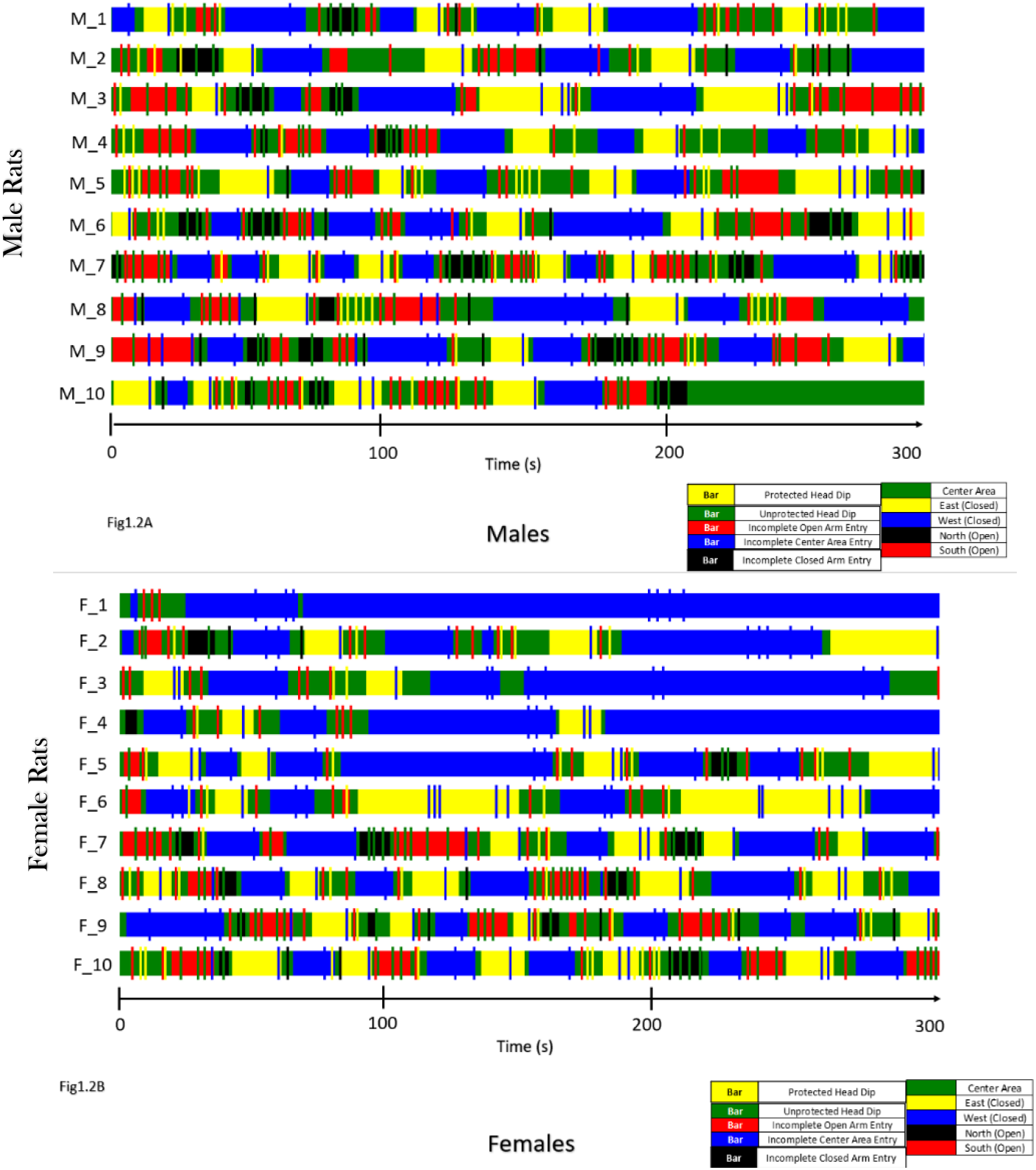
Visual Maps obtained by the Kinoscope software showing the chronological changes in the behaviour. The changes in the behaviour analysed with time in both the groups, males (1.2A) and females (1.2B).

To graphically represent the data obtained by the visual maps, changes observed in the maps were plotted. The behaviour in the open and closed sub-arms was investigated separately according to the design of the EPM and a similar trend was observed with evident preference to one of the two arms (Fig1.3). Although males were observed to spend more time in either of the Open arms, both males (t=−3.6; p=0.002; Cohen’s d=1.45; df=9) and females (t=−1.36; p=0.09; Cohen’s d=3.49; df=9) preferred the Open Arm 2 over Open Arm 1, with males showing a statistically significant difference whereas females showed a trend. In case of the closed arms, both males (t=3.18; p=0.003; Cohen’s d=1.42; df=9) and females (t=3.52; p=0.001; Cohen’s d=1.57; df=9) showed a significant preference to the Closed Arm 1, which was placed on the left side (from the researcher’s viewpoint). Refer to Tables 2 and 3 in the Supplementary Information for more statistical measures.

**Figure 1.3.**
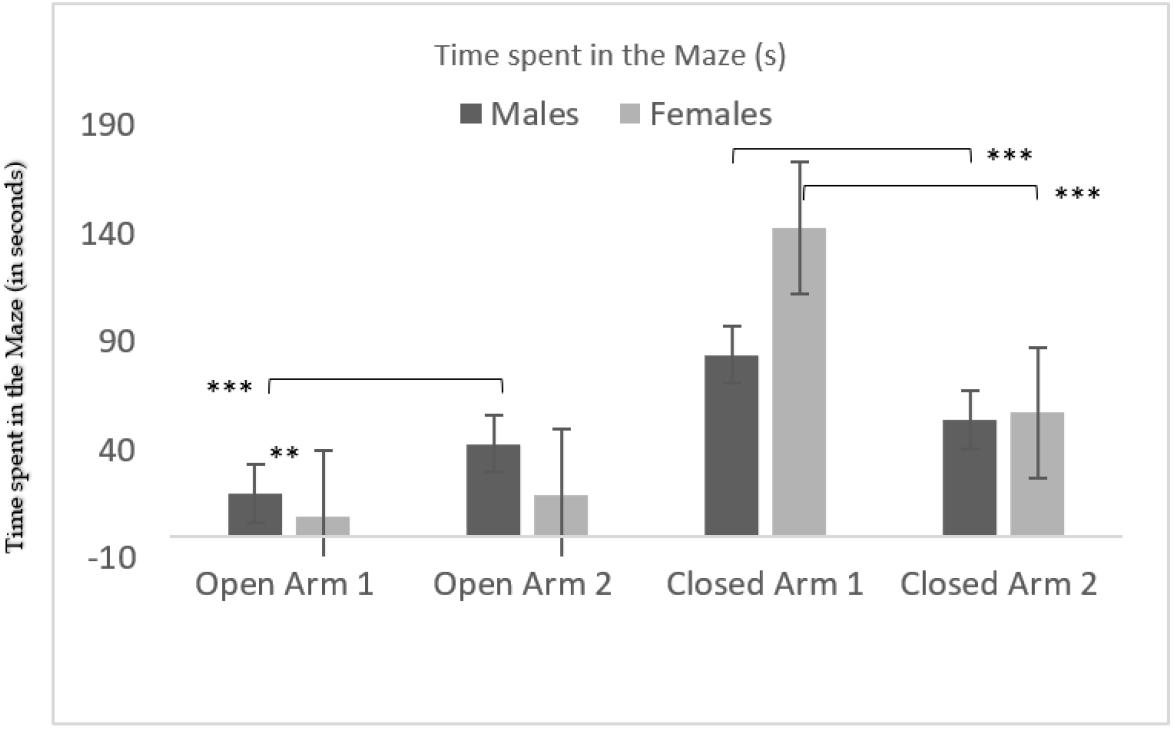
Preference to one of the sub-arms of the Open and Closed arms by both the groups. The figure shows the time spent in each of the sub-arms, Open Arm 1 and 2 and the Closed Arm 1 and 2. A preference to Open Arm 1 and Closed arm 2 was shown by both the groups. (***: p-vlaue<0.05).

### 2. Temporal Feature: Chronology of time spent

Further, to examine how the anxious phenotype of spending less time in the open arms and more time in the closed arms changes within the trial duration of 300s, we time-tagged this particular behaviour. Time spent in different regions within the bins of 50 seconds was extracted by the Kinoscope algorithm. Males showed more time spent in the open arms when compared to the females in all the 6 bins extracted (Fig2.1A) with significant differences in the time bins, 0-50s (t=1.767; p=0.047; Cohen’s d=0.79; df=9) and 50.1-100s (t=2.889; p=0.004; Cohen’s d=1.292; df=9), i.e., initial phase of the trial and 250.1-300s (t=1.868; p=0.039; Cohen’s d=0.835; df=9), i.e., the last 50s of the trial. In case of closed arms however (Fig2.1B), the completely opposite trend was observed with females spending more time in the closed arms as compared to the males with the significant differences in five out of the 6 bins examined; 0-50s (t=−1.95; p=0.033; Cohen’s d=0.873; df=9); 50.1-100s (t=−2.563; p=0.01; Cohen’s d= 1.146; df=9); 100.1-150s (t=−2.114; p=0.024; Cohen’s d=1.146; df=9); 200.1-250s (t=− 2.799; p=0.006; Cohen’s d=1.252; df=9) and 250.1-300s (t=−3.094; p=0.003; Cohen’s d=1.384;df=9). Refer Tables 4 and 5 (Supplementary Information) for the means and other statistical parameters.

**Figure 2.1.**
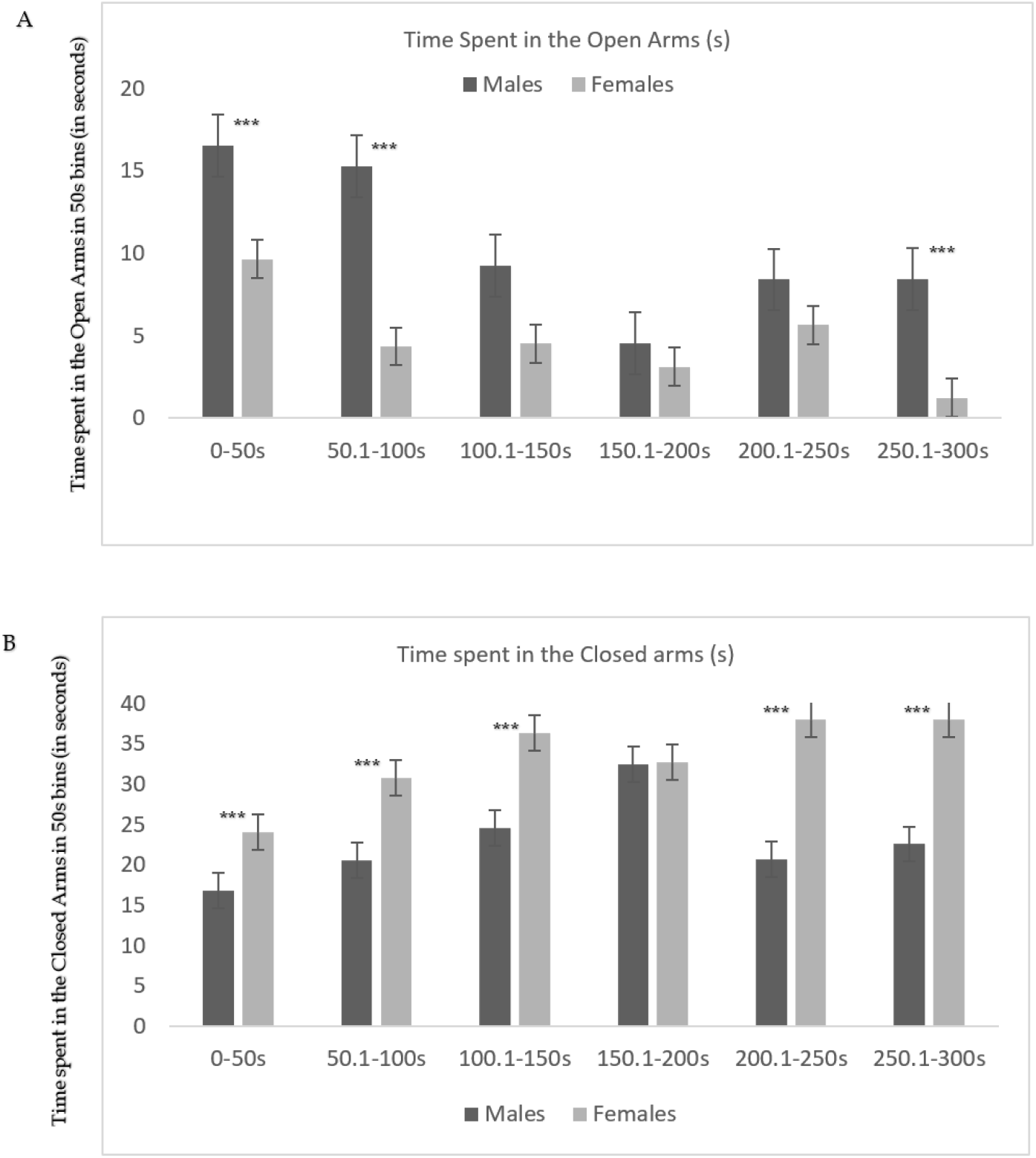
Chronology of the time spent in Open and Closed arms in 50s bins. The trial duration of 300s has been divided into 6 bins of 50s each. Males were shown to spend more time in the open arms in all the bins with significant difference in the initial and last bin (2.1A). Females were shown to spend more time in the closed arms in all the bins with significant differences in 5 out of the 6bins. N=10/group.

#### 2.2. Change in Arm Preference

Next, we examined whether the duration of the time spent in open and closed arm varied across the first and last half of trial duration [i.e., time spent in two blocks of the trial, first block, i.e., first half (0-150s) vs last block, i.e., last half (150-300s) as two within-subject differences between males and females (between-subject variable) (Fig2.2). Result of a 2(male vs female) ×2 (open vs. closed arm) ×2 time (first block vs. last block) mixed analysis of variance showed main effect of open arm F (1, 18) = 94.13, p = .000 partial eta squared = .839, time spent in open arm increased from first to last block (mean 1 = 22.71, mean 2 = 84.38). Interaction of open arm time × sex was significant F (1, 18) = 14.28, p = .001 partial eta squared = .442, time spent in open arm showed greater increase in females (mean 1 = 14.24, mean 2= 99.35) compared to that of males (mean 1 = 31.19, mean 2 = 68.83). Main effect of closed arm was non-significant F (1, 18) = .079, p = .782. Interaction of closed arm × sex was also non-significant F (1, 18) = 1.84, p = .192. Interaction of open arm × closed arm was significant F (1, 18) = 11.17, p = .004, partial eta squared = .383, time spent in open arm declines from first to last half of the trial (mean 1 = 29.76, mean 2 = 15.67) whereas the time spent in closed arm showed an increase from the first to the last half of the trial (mean 1 = 76.56, mean 2 = 92.20). Three-way interaction of open arm × closed arm × sex was non-significant F (1, 18) = .171, p = .684, suggesting that the increase in time spent in closed arm and decrease in time spent in open arm was independent of the sex. Equivalence of variance and normality of data showed no concern.

**Figure 2.2.**
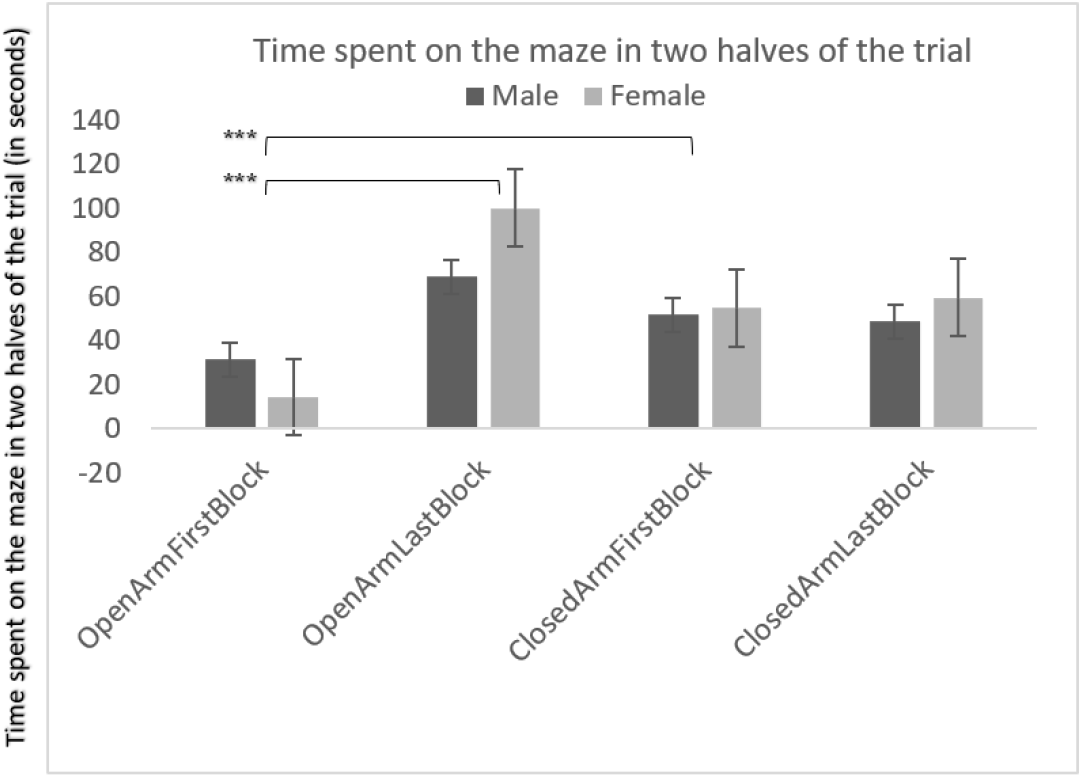
Repeated measures Analysis of the time spent in the Open and Closed arms in the two halves of the trial. The figure shows the increase in time spent in the open arm in the latter half of the trial. This increase was more for the females as compared to the males. N=10/group. Data expressed as mean ± SEM

## Discussion

We examined sex-specific spatial (arm preference) and temporal feature (time bins) in elevated plus maze performance. Results confirmed that males spent more time in the open arms compared to the closed arms, aligned with time spent in the open arms reflecting low anxiety (Walf, 2007; Masse, 2007; Haller, 2012) and increased risk-taking in males (Kessler et al, 2012; Altemus et al, 2014).

Spatial analysis of arm preference, that is two open and closed arms analysed separately indicated a preference for one of the sub-arms in case of both open and closed arms. This sub-arm preference is less documented but observed by some studies reporting side-bias in arm preference (Sherman et al. 1980; Castellano, 1987; Schwarting, 2005). Interestingly, side-bias in arm preference emerged in a sex-specific manner, male spatial preference for one sub-arm was observed in open arms, and female spatial preference for sub-arm was observed in closed arms.

Although the conventional EPM analysis uses average time spent in both the open arms and in the two closed arms, our results indicate males might have specific preference for one of the open arms, and female might show specific preference for one of the closed arms, assuming an indifference in preference for open and closed sub-arms might contribute to mixed, or variable findings for sex-differences in EPM performance. Sex-differences in sub-arm preference for open and closed arm potentially indicated the utility of delineating sub-arm preference.

Similarly, analysis of the time spent across separate time bins helped understand temporal alteration in arm preference, specifically learning across trials that reflects regulation of anxiety and tolerance towards risky open arms. Results align with others who examined temporal changes in two trials of EPM and observed that the time spent in the open arm starts decreasing by the 2^nd^ minute (120s) (Rodgers et al., 1996). Results indicated that males spent more time in the risky open arm in the first two, and the last time segments, indicating that the time spent in open arms as a marker of low anxiety and high risk taking might not be uniform in males. These results align with other reports of non-uniformity in time spent in open arm, for instance, reduced anxiety towards open-arm was observed due to earlier exposure and habituation on comparing two EPM trials, (Schrader et.al, 2018), and even a single-trial showed tolerance towards open-arm after 2 minutes of the EPM trial (Arabo et.al, 2014). Males showed variation in time spent in open arm, however, females showed more consistency, spending more time in the safe closed arm in all the time segments except for the segment 150 – 200 sec.

Further, temporal analysis of time spent in 6 bins showed a decrease in the time spent in the open arms till 2 minutes by both males and females, interestingly, only males showed increased time spent in open arms at the end of the trial. Time spent in the closed arms starts increasing by the 2^nd^ minute for both the sexes, however males showed a decline whereas females show an increase with trials. These findings align with others indicating females at risk of high anxiety (de Visser et al., 2011; Kessler et al, 2012; Altemus et al, 2014; SAMHSA, 2018, Mamlouk et al., 2020 and Bangasser and Cuarenta, 2021). However, temporal analysis of trial duration divided into two halves indicated that larger increase in the time spent in the open arms for females compared to the males, reflecting larger increase in the time spent in open arms, possibly indicative of higher anxiety-regulation compared to the males. Temporal analysis of time spent divided in smaller discrete time bins compared to the commonly used EPM measure of total time spent in open arms for the 5 minutes trial duration might provide a nuanced insight to sex-specific anxiety and risky preference of open arm. Our results indicate the possibility of high anxiety regulation in females and increase in risk-taking, our results corroborate with the rare reports of females exhibiting less anxious behaviour on the EPM (Pavlova et al. 2020; Knight et al., 2021; Bonuti and Murato, 2022).

Despite the widespread use of EPM, and other widely used measures of anxiety, there is a need for improvement in analysing rodent behaviour (Rodgers, 2007), for instance, discriminating between overlapping constructs such as anxiety, risk avoidance, or escape responses (Michalikova, van Rensburg, Chazot, & Ennaceur, 2010), open arm avoidance as learned risk-avoidance rather than an unconditioned risk aversion (Holmes & Rodgers, 1999; Carobrez & Bertoglio, 2005). Detail temporal assessment demonstrates risk taking via initial approach of open arms and closed arms, developing into a distinct closed arm preference for safety (Cole et al., 1996; Holmes & Rodgers, 1998; Rosa et al, 2000; Bertoglio & Carobrez, 2002; Pereira et al. 2005; Casarrubea et al., 2013). This is in contrast to the assumption of unconditioned response (Pellow et al.1985), because approach-avoidance conflict might not be immediately apparent and closed arm preference for safety might be developed with time rather than the default preference (Roy et al., 2009). Spatial and temporal analyses might improve our understanding of underlying neurobiological processes, for instance, to answer questions such as whether the same neural circuitry is manifested differently in males and females, or if different circuities are recruited by males and females to accomplish the same task under anxiety (McCarthy et al., 2012; Becker, 2016; DE Vries, 2004). Some instance, circuitry in anxiety engaged prefrontal-cortex and amygdala (Cahill et al., 2004; Lopez et al., 2011), and frontal-limbic networks show increased connectivity in healthy females than in the males (Lopez, 2011). Preference might change after the initial trials leading to the avoidance of the risky open arms (i.e., less exploration of open arms) until rodents have learnt the spatial features of the maze (Rodgers & Shepherd, 1993; Bertoglio & Carobrez, 2000; Roy et al., 2009; Casarrubea et al., 2013), whether this change in preference differs across male and female rodents might help us understand neurobiological basis of sex-differences in anxiety and risk taking.

## Conclusion

Our understanding of males and females’ anxiety, tolerance and risk preference for open arms in EPM might be improved by adding spatial and temporal analysis of EPM, producing alternate, novel insights such as the possibility of females showing larger increase in open arm time spent, potentially showing greater regulation of anxiety with increased trial exposure.

## Ethical Clearance

The study had an Ethical clearance from the Central Animal Facility, All India Institute of Medical Sciences, New Delhi (Ethical Number – 311/IAEC-1/2022).

## Acknowledgement

We acknowledge the support of Department of Science and Technology – Cognitive Science Research Initiative Grant (DST/CSRI/2018/100(G)) given to the corresponding author, for partially supporting this work as a part of doctoral thesis.

