## Supplementary material for "Examining Anxiety and Risk-taking in Healthy Male and Female Wistar Rats using Spatial and Temporal Analysis of Elevated Plus Maze": Tables 1 - 9

### Supplementary Information

| Time Spent on the Maze (s) | | | | | |
|---|---|---|---|---|---|
| | **Males (Mean±SEM)** | **Females (Mean+SEM)** | **p-values** | **t-values** | **effect size (Cohen's d)** |
| Open Arms | 62.38±2.05 | 28.48±1.87 | 0.005 | 2.87 | 1.29 |
| Closed Arms | 137.68±2.05 | 199.87±1.49 | 0.001 | -3.87 | 1.73 |

**Table 1: Time spent on the maze.** The table shows the time spent in seconds by both groups, Males and Females in the Open and Closed Arms along with other statistical measures. N=10/group. Data was found to have normal distribution.

**Time spent on the Maze, separately in the sub-arms of Open and Closed Arms**

| | **Males (Mean±SEM)** | **Females (Mean+SEM)** | **p-values** | **t-values** | **effect size (Cohen's d)** |
|---|---|---|---|---|---|
| Open Arm 1 | 19.63±2.8 | 9.31±3.3 | 0.025 | 2.11 | 0.94 |
| Open Arm 2 | 42.76±2.3 | 19.17±2.1 | 0.008 | 2.65 | 1.19 |
| Closed Arm 1 | 83.98±1.9 | 142.61±1.2 | 0.009 | -2.58 | 1.16 |
| Closed Arm 2 | 53.70±2.6 | 57.26±1.6 | 0.392 | -0.28 | 0.12 |

**Table 2: Time spent on the maze, in the sub-arms.** The table shows the time spent in seconds by both groups, Males and Females, separately in the sub-arms of the Open and Closed Arms along with other statistical measures. N=10/group. Data was found to have normal distribution.

| | **OA1 (Mean±SEM)** | **OA2 (Mean±SEM)** | **p-value** | **t-values** | **effect size (Cohen's d)** |
|---|---|---|---|---|---|
| Males | 19.63±2.8 | 42.76±2.3 | 0.002 | -3.26 | 1.45 |
| Females | 9.31±3.3 | 19.17±2.1 | 0.09 | -1.36 | 3.49 |
| | **CA1 (Mean±SEM)** | **CA2 (Mean±SEM)** | | | |
| Males | 83.98±1.9 | 53.7±2.6 | 0.003 | 3.18 | 1.42 |
| Females | 142.6±1.2 | 57.26±1.6 | 0.001 | 3.52 | 1.57 |

**Table 3: Time spent on the Maze (in separate sub-arms).** The table shows the time spent separately in the sub-arms, Open Arm 1 (OA1), Open Arm 2 (OA2), Closed Arm 1 (CA1) and Closed Arm 2 (CA2), along with other statistical measures. N=10/group. Data was found to have normal distribution.

**Chronology of the Time Spent in the Open Arms (s)**

| | Males (Mean±SEM) | Females (Mean+SEM) | p-value | t-value | cohen's d (Effect Size) |
|---|---|---|---|---|---|
| 0-50s | 16.51±3.28 | 9.64±3.52 | 0.047 | 1.767 | 0.790 |
| 50.1-100s | 15.27±3.26 | 4.34±3.69 | 0.004 | 2.889 | 1.292 |
| 100.1-150s | 9.25±3.41 | 4.51±3.42 | 0.116 | 1.239 | 0.554 |
| 150.1-200s | 4.52±3.71 | 3.11±3.71 | 0.335 | 0.434 | 0.194 |
| 200.1-250s | 8.41±3.37 | 5.65±3.54 | 0.236 | 0.734 | 0.328 |
| 250.1-300s | 8.43±2.94 | 1.23±5.06 | 0.039 | 1.868 | 0.835 |

**Table 4: Chronology of the time spent in the Open Arms.** The table shows the chronology of the time spent in the open arms in the bins of 50 seconds each, along with the other statistical measures. N=10/group. Data was found to have normal distribution.

**Chronology of the Time Spent in the Closed Arms (s)**

| | Males (Mean±SEM) | Females (Mean±SEM) | p-value | t-value | cohen's d (effect size) |
|---|---|---|---|---|---|
| 0-50s | 16.82±3.78 | 24.08±3.25 | 0.033 | -1.951 | 0.873 |
| 50.1-100s | 20.58±3.50 | 30.82±3.22 | 0.010 | -2.563 | 1.146 |
| 100.1-150s | 24.57±2.94 | 36.27±2.76 | 0.024 | -2.114 | 0.945 |
| 150.1-200s | 32.44±3.39 | 32.67±2.72 | 0.482 | -0.047 | 0.021 |
| 200.1-250s | 20.69±2.68 | 38.05±2.69 | 0.006 | -2.799 | 1.252 |
| 250.1-300s | 22.58±2.84 | 37.98±3.21 | 0.003 | -3.094 | 1.384 |

**Table 5: Chronology of the time spent in the Closed Arms.** The table shows the chronology of the time spent in the closed arms in the bins of 50 seconds each, along with the other statistical measures. N=10/group. Data was found to have normal distribution.

**Chronology of the time spent in the sub-arm Open Arm 1**

| | Males Mean±SEM | Females Mean+SEM | t-value | p-value | cohen's d (effect size) |
|---|---|---|---|---|---|
| 0-50s | 3.18±4.39 | 3.33±5.38 | -0.07309 | 0.47127 | 0.032688 |
| 50-100s | 7.68±3.89 | 1.46±5.33 | 2.63156 | 0.008468 | 1.176869 |
| 100-150s | 2.27±4.39 | 0.01 | 1.37352 | 0.093229 | 0.614257 |
| 150-200s | 1.71±4.29 | 1.43±5.74 | 0.14644 | 0.442601 | 0.065489 |
| 200-250s | 2.17±4.65 | 3.10±4.46 | -0.4343 | 0.334617 | 0.194223 |
| 250-300s | 2.61±4.34 | 0.01 | 1.54737 | 0.069588 | 0.573419 |

**Table 6: Chronology of the time spent in the sub-arm Open Arm 1.** The table shows the chronology of the time spent in the sub-arm Open Arm 1 in the bins of 50 seconds each, along with the other statistical measures. N=10/group. Data was found to have normal distribution.

**Chronology of the time spent in the sub-arm Open Arm 2**

| | **Males** | **Females** | **t-value** | **p-value** | **cohen's d** |
|---|---|---|---|---|---|
| | **Mean±SEM** | **Mean±SEM** | | | **(effect size)** |
| 0-50s | 13.33±3.13 | 6.32±4.16 | 1.891 | 0.037 | 0.846 |
| 50-100s | 7.59±4.29 | 2.89±4.66 | 2.086 | 0.026 | 2.073 |
| 100-150s | 6.98±3.84 | 4.51±3.42 | 0.718 | 0.241 | 0.321 |
| 150-200s | 2.80±4.72 | 1.68±4.74 | 0.562 | 0.291 | 0.251 |
| 200-250s | 6.24±3.57 | 2.54±4.31 | 1.230 | 0.117 | 0.678 |
| 250-300s | 5.81±2.92 | 1.23±5.06 | 1.175 | 0.128 | 0.526 |

**Table 7: Chronology of the time spent in the sub-arm Open Arm 2.** The table shows the chronology of the time spent in the sub-arm Open Arm 2 in the bins of 50 seconds each, along with the other statistical measures. N=10/group. Data was found to have normal distribution.

**Chronology of the time spent in the sub-arm Closed Arm 1**

| | **Males** | **Females** | **t-value** | **p-value** | **cohen's d** |
|---|---|---|---|---|---|
| | **Mean±SEM** | **Mean+SEM** | | | **(effect size)** |
| 0-50s | 10.03±3.61 | 16.27±3.12 | -1.535 | 0.071 | 0.686 |
| 50-100s | 13.30±3.43 | 21.97±2.72 | -1.716 | 0.052 | 0.767 |
| 100-150s | 16.02±3.1 | 25.51±2.27 | -1.366 | 0.094 | 0.611 |
| 150-200s | 20.85±3.18 | 23.91±2.52 | -0.521 | 0.304 | 0.233 |
| 200-250s | 12.08±3.71 | 30.96±2.31 | -2.968 | 0.004 | 1.327 |
| 250-300s | 11.69±2.84 | 23.99±2.45 | -1.870 | 0.039 | 0.836 |

**Table 8: Chronology of the time spent in the sub-arm Closed Arm 1.** The table shows the chronology of the time spent in the sub-arm Closed Arm 1 in the bins of 50 seconds each, along with the other statistical measures. N=10/group. Data was found to have normal distribution.

**Chronology of the time spent in the sub-arm Closed Arm 2**

| | **Males** | **Females** | **t-value** | **p-value** | **cohen's d** |
|---|---|---|---|---|---|
| | **Mean±SEM** | **Mean+SEM** | | | |
| 0-50s | 6.78±3.72 | 7.81±3.65 | -0.312 | 0.379 | 0.139 |
| 50-100s | 7.28±3.33 | 8.85±3.72 | -0.430 | 0.336 | 0.192 |
| 100-150s | 8.55±4.27 | 10.76±2.65 | -0.457 | 0.327 | 0.204 |
| 150-200s | 11.59±3.57 | 8.76±3.61 | 0.814 | 0.213 | 0.364 |
| 200-250s | 8.61±3.1 | 7.09±2.68 | 0.278 | 0.392 | 0.124 |
| 250-300s | 10.89±3.06 | 13.99±2.76 | -0.580 | 0.285 | 0.259 |

**Table 9: Chronology of the time spent in the sub-arm Closed Arm 2.** The table shows the chronology of the time spent in the sub-arm Closed Arm 2 in the bins of 50 seconds each, along with the other statistical measures. N=10/group. Data was found to have normal distribution.

**Chronology of the ratio of the time spent in the Open Arms / Closed Arms**

| | **Males (Mean+SEM)** | **Females (Mean+SEM)** | **t-value** | **p-value** |
|---|---|---|---|---|
| 0-50s | 1.27±10.68 | 0.66±11.4 | 1.669 | 0.056 |
| 50.1-100s | 1.26±7.4 | 0.21±16.03 | 1.779 | 0.046 |
| 100.1-150s | 0.67±11.36 | 0.30±11.84 | 1.108 | 0.141 |
| 150.1-200s | 0.18±17.7 | 0.26±12.43 | -0.322 | 0.376 |
| 200.1-250s | 0.67±9.28 | 0.29±14.43 | 0.951 | 0.177 |
| 250.1-300s | 0.75±16.69 | 0.04±27.13 | 1.699 | 0.053 |

**Table 10: Chronology of the ratio of the time spent in the Open Arms/Closed Arms.** The table shows the chronology of the ratio of the time spent in the open arms/closed arms in the bins of 50 seconds each, along with the other statistical measures. N=10/group. Data was found to have normal distribution.
